# Differences in T-cell counts and neighborhood patterns in human colorectal adenomas and sessile serrated lesions

**DOI:** 10.64898/2026.01.06.697786

**Authors:** Souvik Seal, Lauren R. Fanning, Evan Bagley, Christine Bookhout, Elizabeth L. Barry, Elizabeth O’Quinn, Dale C. Snover, David N. Lewin, Silvia Guglietta, Antonis Kourtidis, Martha J. Shrubsole, John A. Baron, Todd A. Mackenzie, Alexander V. Alekseyenko, Kristin Wallace

**Affiliations:** Hollings Cancer Center, Medical University of South Carolina, Charleston, SC, USA; Department of Public Health Sciences, Medical University of South Carolina, Charleston, SC, USA; Department of Pathology and Laboratory Medicine, University of North Carolina School of Medicine, Chapel Hill, NC, USA; Department of Epidemiology, Geisel School of Medicine at Dartmouth, Lebanon, NH, USA; Biorepository & Tissue Analysis Shared Resource, Hollings Cancer Center, Medical University of South Carolina, Charleston, SC, USA; Department of Pathology, Fairview Southdale Hospital, Edina, MN, USA; Department of Pathology and Laboratory Medicine, Medical University of South Carolina, Charleston, SC, USA; Department of Regenerative Medicine and Cell Biology, Medical University of South Carolina, Charleston, SC, USA; Department of Medicine, Vanderbilt-Ingram Cancer Center, Nashville, TN, USA; Department of Epidemiology, University of North Carolina School of Medicine, Chapel Hill, NC, USA; Department of Biomedical Data Science, Geisel School of Medicine at Dartmouth, Lebanon, NH, USA

## Abstract

**Background:** T-cell responses influence recurrence and survival in colorectal cancer. T-cell subset distributions vary by molecular phenotype and location, shaping cytotoxic or immune-cold tumor immune microenvironments (TiMEs). However, the T-cell contexture and their spatial proximities within preinvasive lesions are not well characterized.

**Methods:** We analyzed sessile serrated lesions (SSLs), tubulovillous/villous adenomas (TVs), and tubular adenomas (TAs) from three completed studies (N=120). Whole-slide multiplex immunofluorescence was used to quantify eight T-cell subsets (CD4⁺, CD8⁺, Th1, Th17, Treg, Tc1, Tc17, TcTreg). Counts were compared by histology using a generalized linear mixed model with a negative binomial distribution, including an offset for total cell counts and adjusting for age, sex, anatomic location, and lesion size. Nearest-neighbor (NN) analyses assessed proximities of T-cell pairs across lesion types.

**Results:** TAs and SSLs had higher CD4⁺ and CD8⁺ T-cell counts compared with TVs (q<0.05). SSLs had lower Th17 counts than TAs (q<0.05) and compared with TVs leaned toward fewer Tregs (q=0.07). NN analysis showed that TVs, compared with SSLs and TAs, had increased Treg clustering. In contrast, TA versus SSL comparisons revealed predominant CD4⁺ clustering with Th17, Th1, and CD8⁺ subsets.

**Conclusion:** TVs are characterized by lower T-cell infiltration and a greater tendency for regulatory T-cell clustering, consistent with an immune-cold TiME relative to TAs. SSLs and TAs were both more immune-infiltrated than TVs, but SSLs appeared less inflamed and less dominated by regulatory subsets. In contrast, CD4⁺dominant clustering in TAs suggested stronger helper coordination. Preinvasive lesions therefore demonstrate immune and spatial heterogeneity by lesion types.

## Introduction

T-cell infiltration is a strong predictor of recurrence and survival in colorectal cancer (CRC). ^1–4^ The distribution of helper T-cell subsets varies by tumor molecular phenotype and anatomic site. ^5–7,8^ Microsatellite instability–high (MSI–H) CRCs, which arise predominantly in the proximal colon,^9, 10^ are enriched for CD8⁺ cytotoxic T-cells and Th1 cells, whereas microsatellite-stable (MSS) tumors, which are more common in the distal colorectum,^7, 11^ tend to have higher Th17 cell densities. Prior studies have shown better CRC outcomes for greater infiltration of Th1 and CD8^+^ cytotoxic T-cells.^1–4^

Patterns of T-cell infiltration are characterized far less in preinvasive lesions than in CRCs. Sessile serrated lesions (SSLs), which are predominately located in the proximal colon and resemble MSI–H CRCs genomically and immunologically, are dense with intraepithelial lymphocytes including regulatory T-cells (Tregs) and CD4^+^ memory T-cells (CD4^+^CD45RA^-^CD45RO^+^);^6, 9, 12, 13^ tubular adenomas (TAs), part of the canonical MSS pathway, occur throughout the colorectum and often show Th1- and Th17-associated cytokine signatures;^5^ and tubulovillous/villous adenomas (TVs), frequently located in the distal colon and likely related to *KRAS* mutated MSS CMS3 CRCs, exhibit immune-cold profiles that intensify with increasing villous architecture.^7, 10, 13^

Beyond differences in average counts, spatial proximity among T-cell subsets may shape immune function within the tumor microenvironment (TME).^14, 15^ T-cells cluster with other T-cells to form niches where close contact and reciprocal (paracrine) signaling help to sustain T-cell activation and function:^16,17^ specific neighborhood relationships may reinforce cytotoxic-effector (Tc1/Th1) interactions versus immunoregulatory (Th17/Treg) milieus, whereas sparse or lack of T-cell neighbors may mark immune-cold regions or “immune deserts.”^18^ Although few studies have examined T-cell proximities in CRCs, Huang and colleagues^14^ reported strong Tc1-Th1 spatial co-localization (also called interactions) in MSI-H CRCs and conversely, higher Th17, Treg, and Th2 interactions in MSS tumors. In prior pilot work,^19^ we observed distinct spatial relationships in preinvasive lesions: SSLs contained abundant CD4⁺ T-cells but Tregs were sparse and spatially distant from other CD4^+^ T-cells, whereas conventional adenomas showed frequent CD4^+^/Treg commingling, consistent with a coordinated immune dampening response. However, to our knowledge, no study has compared T-cell neighbor relationships between different histological types of preinvasive lesions.

Therefore, in the present study, we used single-cell multiplex immunofluorescence (mIF) analysis to profile eight helper and cytotoxic T-cell subsets by histologic type (TA, TV, and SSL) adjusting for anatomic location, age, sex and lesion size using a convenience sample drawn from a larger multi-center retrospective study. We then examined pairwise spatial relationships among eight T-cell subsets by histologic type using a nearest neighbor (NN) approach.^20, 21^

## Methods

Our study is a cross-sectional analysis which compares T-cell counts across three lesion types. Samples came from participants in three completed studies: the Vitamin D/Calcium Polyp Prevention Study (ViCAPPS, 2004-2013),^22^ Diet and Health Studies III–V (DHS, 1998–2010),^23,24^ and the Tennessee Colorectal Polyp Study (TCPS, 2003–2015).^25^ In each study, participants had been diagnosed with one or more histologically confirmed colorectal adenomas or SSLs. Exclusion criteria included: familial CRC syndromes, inflammatory bowel disease, prior colon resection, and history of cancer other than non-melanoma skin cancer within at least 5 years prior to enrollment.

Each clinical study (ViCAPPS, DHS III-V, TCPS) provided four unstained 5 µm FFPE sections per subject. Prior to shipment, a site pathologist verified that the block contained ≥5 mm of tissue, was colonic in origin, and had adenomatous or serrated histology. Slides were barcoded, deidentified, and shipped in batches to the Data Coordinating Center at the Medical University of South Carolina (MUSC). Upon receipt all samples were verified against manifests by the pathology coordinator and stored in a desiccator in a cold room until multiplex immunofluorescence (mIF) analysis was started.

To ensure standardization of diagnosis across the study samples, a central pathology review was performed by an expert in gastrointestinal pathology (D.S.) blinded to original diagnosis and clinical information. For each digitized H&E, the pathologist identified the histologic type (e.g., TA, TV, SSL) and The diagnosis was recorded within a REDCap database. If there was discordance between the study pathologist and original diagnoses regarding histologic type of adenoma (TA, TV), the study pathologist’s diagnosis was accepted. If there was a disagreement between the study pathologist and the original diagnosis on whether the lesion was adenomatous, or serrated in origin, a second study gastrointestinal pathologist (D.L.) performed an independent review on the same digitized slide and was blinded to the first study pathologist diagnosis. To resolve differences between the first and second study pathologists, an adjudication protocol was used, and disagreements were resolved by reviewing images together and arriving at a consensus. Results presented here reflect the consensus diagnosis.

To reduce potential variability across mIF runs, specimens were classified into four groups to ensure the groups were equally distributed across mIF batches: small (<10 mm) TAs, large (≥10 mm) TAs, TVs, and SSLs. Each 24-slide mIF batch contained 22 study samples balanced by clinical study (ViCAPPS, DHS, TCPS) and each of the four strata, plus two control tissues (normal colon mucosa and tonsil). Antibody panels and staining order were optimized using tonsil control, normal control and 20 colorectal neoplasms (10 adenomas (½ TAs, ½ TVs), 10 SSLs) balanced by anatomic location (proximal, distal, rectal). Control tissues for optimization were obtained from a commercial source or MUSC surgical pathology archives (normal control, adenomas, and SSLs) with aid from study pathologist (D.L.).

Staining was performed on the Roche Ventana Discovery Ultra platform with Akoya Opal™ tyramide signal amplification, following methods previously described.^26^ Slides were stained for five immune markers (CD4, T-bet (TBX21), FoxP3, RORγt, CD8) and Pan Keratin, followed by DAPI nuclear counterstain for cell detection (see Supplemental Table 1 for additional details). Stained slides were mounted with ProLong™ Gold Antifade reagent, coverslipped and imaged at 20× magnification on the Akoya Vectra® Polaris™ (PhenoImager™ HT). Whole-slide images were reviewed and annotated in Phenochart™ by staff blinded to the diagnosis.

Image analysis was conducted in inForm® Tissue Analysis Software (Akoya Biosciences), with downstream processing in the Phenoptr Reports R package. Tissue segmentation, cell segmentation, and phenotyping were trained on representative regions from each lesion type in the pilot study (see Supplementary Methods for complete description). T-cell phenotypes included CD4⁺, Th1 (CD4^+^T-bet^+^), Th17 (CD4^+^RORγt^+^), Treg (CD4^+^FoxP3^+^), CD8⁺, Tc1 (CD8⁺T-bet^+^), Tc17 (CD8^+^RORγt^+^), and TcTreg (CD8^+^FoxP3^+^). DAPI stain, a nuclear marker, was used to define individual cells. Pan-keratin was selected to aid in epithelial membrane detection (keratin positive) to distinguish epithelial and stromal tissue regions.

Cells negative for all T-cell markers were classified as “Other” (i.e., negative for CD4/CD8/T-bet/RORγt/FoxP3, positive or negative for Pan-keratin). Although these cells lack specific biological identity within the context of the current study, their presence, when occurring in contiguous regions, could indicate regions suspicious for T-cell deserts.^7^ That said, these T-cell sparse regions likely still contain other immune cell types or other markers we did not measure (e.g., macrophages, dendritic cells, collagens, mucins). Details of image analysis training, segmentation parameters, and phenotyping are provided in the Supplementary Methods.

## Statistical Methods

For each T-cell phenotype, we assessed differences in whole slide counts across three histologic types (TA, TV, SSL). Because cell counts are often overdispersed (i.e., variance exceeds that expected by the binomial distribution), we fit a generalized linear mixed model with a negative binomial (NB) distribution and log link. To account for variable tissue area/cellularity in different lesions we included the log of the total cell count (including T-cell and “Other” cells) as an offset in the model generating “normalized” mean counts. To avoid confounding, the model was adjusted for fixed effects of age, sex, anatomic location, and lesion size. To mitigate potential batch effects, clinical study and data-collection batch were adjusted for as random effects. In each model, we generated covariate-adjusted mean T-cell counts for each immune cell type and their 95% CIs for each histologic type using estimated marginal means (emmeans) statement in R. We also computed the rate ratios (RR) for each T-cell type for each lesion pair (i.e., SSL vs. TA, SSL vs. TV, TA vs. TV) at a false discovery rate (FDR) controlled at q< 0.05.

To assess spatial co-localization between pairs of T-cells, we performed a neighborhood analysis by integrating established methods.^20, 21, 27^ For each lesion, a specific T-cell type A (the “anchor”) was selected, and for each of that type of T-cell, the nearest T-cell type B (its nearest neighbor (NN)) was recorded (**Supplemental Figure 1**). We then computed the proportion of these neighbors that were of type B (i.e., “NN”). For anchors and neighbors, we considered all previously defined cell types, i.e., CD4^+^, CD8^+^, Th1, Th17, Tc1, Tc17, Treg, TcTreg. This observed proportion was adjusted by the expected proportion under the null hypothesis of complete spatial randomness, ^28, 29^ estimated via a bootstrap approach that permuted cell labels 50 times. (**Supplemental Figure 1**) It should be clarified that such an adjustment is crucial to ensure that the NN results are not driven simply by cell type abundance or underlying tissue deformities^29^. The adjusted proportions were averaged within each subject, and group-level differences across histologic types were assessed using the Kruskal-Wallis test to detect differential directed interactions (A → B). We also completed a NN analysis using closest distance to the “Other” cell types. Here we also compared the three histologic types as above but did not permute an expected proportion under the null.

## Data availability

The data generated in this study are available upon request from the corresponding author.

## Results

We analyzed samples from 120 participants (**Table 1**). Lesion histologic types differed by sex, clinical study, lesion size, and anatomic location. Women had a higher proportion of TVs than men. TVs occurred more often in the distal colon and rectum and were larger, on average, than SSLs and TAs. Six traditional serrated adenomas (TSAs) were excluded from analysis due to small sample size.

**Table 1.**
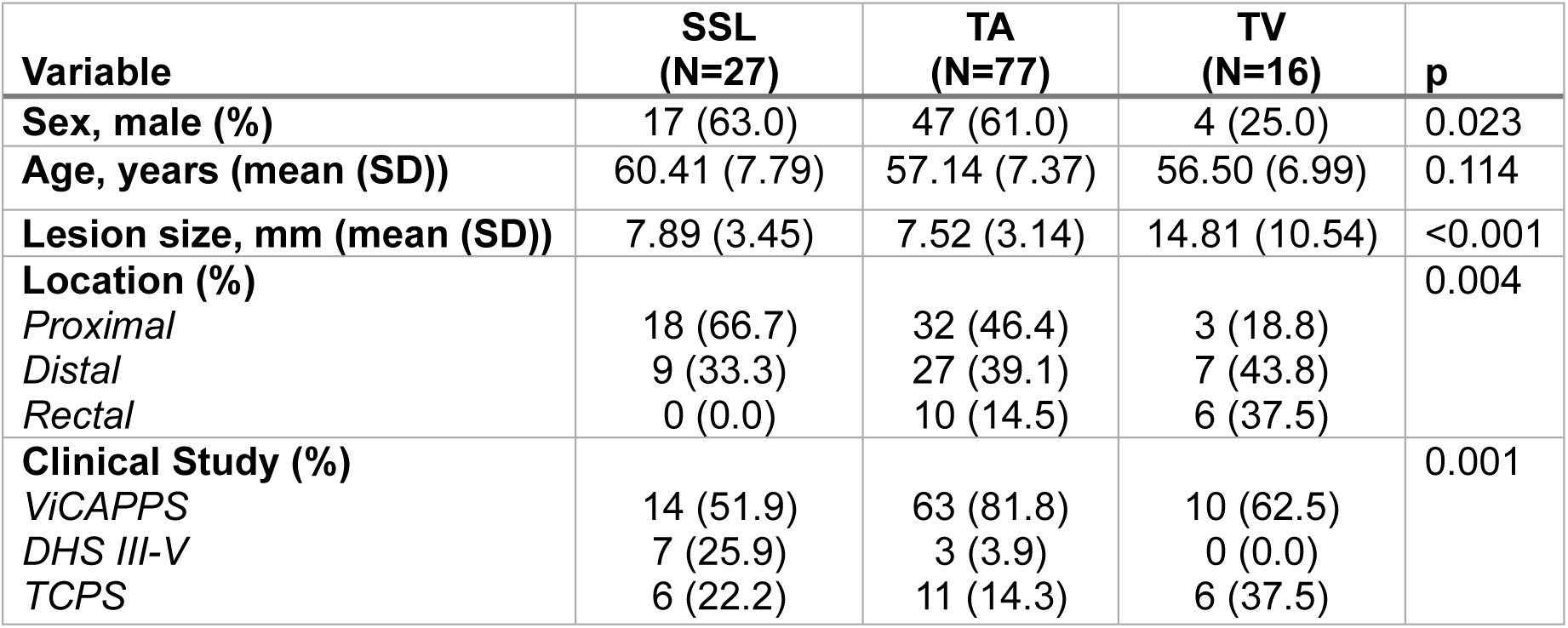
Distribution of patient, lesion, and site variables across lesion histologic types (SSL, TA, TV). *ViCAPPS*: Vitamin D/Calcium Polyp Prevention Study; *DHS III-V*: Diet and Health Studies III-V; *TCPS*: Tennessee Colorectal Polyp Study

Mean T-cell counts and their RRs are shown in **Table 2**, **Supplemental Figure 2, and Supplemental Table 2**. Compared with TVs, TAs had higher counts of both helper and cytotoxic T cells: CD4⁺ (TA: TV ratio = 2.07, q<0.001) and CD8⁺ (TA:TV 1.54, q=0.03). Th17 counts were also higher in TAs than in TVs (TA:TV ratio = 2.04, q=0.004). SSLs showed a similar pattern versus TVs, with higher CD4⁺ (SSL:TV ratio = 1.85, q=0.004) and CD8⁺ (SSL:TV ratio 1.82, q=0.007). In contrast, Tregs and TcTregs were enriched in TVs, though did not retain statistical significance after correcting for multiple comparisons: the SSL:TV Treg and TcTreg ratios were 0.41 (q=0.07) and 0.36 (q=0.09), respectively. No statistically significant differences were observed between SSLs and TAs, although SSLs showed a nonsignificant tendency toward lower Th1 counts relative to TAs (0.54, q=0.19) (**Supplemental Figure 2**)

**Table 2.** Mean T-cell counts, and 95% CIs in SSLs, TAs, and TVs normalized by lesion total cells counts and adjusted for age, sex, anatomic location and corrected at q < 0.05.

| Cell Type | SSL Count (95% CI) | TA Count (95% CI) | TV Count (95% CI) |
| --- | --- | --- | --- |
| **CD4** | 23,629 (19,132-29,184) | 26,944 (23,689-30,646) | 13,129 (9,923-17,365) |
| **CD8** | 3,808 (3,048-4,758) | 3,333 (2,896-3,837) | 2,082 (1,542-2,810) |
| **Th1** | 216 (104-448) | 429 (233-792) | 286 (125-657) |
| **Th17** | 1,654 (1,189-2,302) | 2,156 (1,773-2,623) | 1,090 (720-1,651) |
| **Treg** | 286 (176-465) | 406 (303-545) | 781 (420-1,449) |
| **Tc1** | 159 (86-296) | 222 (131-375) | 203 (101-409) |
| **Tc17** | 302 (209-436) | 306 (244-383) | 218 (138-343) |
| **TcTreg** | 9 (5-18) | 13 (8-19) | 28 (13-61) |

In the neighborhood analyses comparing TAs vs TVs, we identified five statistically significant anchor-NN pairs (q<0.05).(**Table 3**; **Supplemental Figure 3**). Four of five pairs involved a regulatory phenotype (Treg or TcTreg) as the nearest neighbor. The most abundant anchor-NN pairs were CD4^+^/TcTreg, CD8^+^/TcTreg, CD4^+^/Treg, TcTreg/CD8^+^, and Th17/Treg. These results point to Treg and TcTreg as more common neighbors in TVs relative to TAs. Representative images (**Supplemental Figure 4**) illustrate tendency for TVs relative to TAs to exhibit closer proximities between Tregs and other helper T-cells.

**Table 3.** Nearest neighbor analysis: Anchor and neighbor cell types, proportions (%), and q-values for TA vs. TV , SSL vs. TV, and SSL vs. TA comparisons.

| TA vs. TV |  |  |  |  |
| --- | --- | --- | --- | --- |
| Anchor Cell Type | NN Cell Type | TA (%) | TV (%) | q-value |
| CD4<sup>+</sup> | TcTreg | 0.004 | 0.04 | 0.003 |
| CD4<sup>+</sup> | Treg | 0.51 | 1.31 | 0.02 |
| Th17 | Treg | 0.64 | 1.46 | 0.02 |
| TcTreg | CD8<sup>+</sup> | 0.98 | 2.29 | 0.007 |
| CD8<sup>+</sup> | TcTreg | 0.04 | 0.15 | 0.05 |
| SSL vs. TV |  |  |  |  |
| Anchor Cell Type | NN Cell Type | SSL (%) | TV (%) | q-value |
| CD4<sup>+</sup> | Treg | 0.32 | 1.31 | 0.004 |
| Treg | Treg | 0.84 | 2.71 | 0.005 |
| **TcTreg** | CD8<sup>+</sup> | 0.86 | 2.29 | 0.01 |
| **Tc17** | Tc17 | 0.75 | 1.66 | 0.03 |
| Th17 | Treg | 0.37 | 1.46 | 0.005 |
| Tc17 | Treg | 0.12 | 0.88 | 0.004 |
| CD8<sup>+</sup> | Treg | 0.06 | 0.69 | 0.004 |
| CD4<sup>+</sup> | TcTreg | 0.01 | 0.04 | 0.004 |
| Treg | Tc17 | 0.04 | 0.35 | 0.04 |
| TcTreg | Treg | 0.04 | 1.09 | 0.007 |
| SSL vs. TA |  |  |  |  |
| **Anchor Cell Type** | **NN Cell Type** | **SSL (%)** | **TA (%)** | **q-value** |
| CD8<sup>+</sup> | CD4<sup>+</sup> | 8.94 | 15.68 | 0.002 |
| CD4<sup>+</sup> | CD4<sup>+</sup> | 21.65 | 28.99 | 0.004 |
| Th17 | CD4<sup>+</sup> | 16.99 | 23.68 | 0.004 |
| Tc17 | CD4<sup>+</sup> | 6.43 | 12.16 | 0.008 |
| **Tc1** | CD4<sup>+</sup> | 4.23 | 7.76 | 0.036 |

Comparing SSLs to TVs, we observed ten significant neighbor pairs (q<0.05; **Supplemental Figure 3**). Seven of the ten anchor-NN pairs involved Treg/TcTreg as the nearest neighbor. The most common pairs were TcTreg/Treg, followed by CD8^+^/Treg, Treg/Tc17, Tc17/Treg, and CD4^+^/TcTreg. (**Table 3**) TVs vs. SSLs also show a tendency to form Tc17-Treg reciprocal pairs, which was not observed in comparison to TAs. Representative images (**Supplemental Figure 4**) illustrate tendency for TVs relative to SSLs to exhibit closer proximities between Tregs and other T-cells, particularly the RORγt^+^ T-cell subtypes (Th17, Tc17).

In contrast to the bulk-count analysis where no significant differences were observed between SSLs and TAs, the neighborhood analysis identified five significant anchor-NN pairs enriched in TAs relative to SSLs (q<0.05; **Table 3**, **Supplemental Figure 3**). These pairs all featured CD4⁺ as the nearest neighbor in TAs relative to SSLs, with Tc17/CD4⁺ as the most prominent pair, followed by Tc1/CD4^+^, CD8^+^/CD4^+^, Th17/CD4^+^, and CD4^+^/CD4^+^.This pattern positions CD4⁺ as the primary neighbor in TAs relative to SSLs, as shown in the representative TA image (**Supplemental Figure 4**).

Lastly, we considered a neighborhood analysis contrasting the abundance of “Other” cells (i.e., non-immune or other immune cells) by histologic type. These analyses were not adjusted for random labeling. “Other” cells appeared as NNs more often in TVs than in TAs and, to a lesser extent, in TVs than in SSLs (**Supplemental Figure 5**). Six of eight T-cell phenotypes as anchor cells had higher “Other” NNs in TVs vs TAs. In SSLs, only Treg, Tc17, and TcTreg as anchor cells showed lower “Other” neighbors relative to TVs. These patterns suggest a relative sparseness of T-cells in TVs relative to both TAs and SSLs. Stroma- and epithelium-specific mean T-cell counts and rate ratios for histologic type comparisons were largely similar to that of the overall tissue analysis and may be found in the Supplemental Material (Supplemental Table 3a-b, Supplemental Table 4a-b).

## Discussion

### TVs as immune colder lesions with T-regulatory and Th17 skewing relative to TAs and SSLs

In prior work,^5^ increasing villous histology in colorectal adenomas has been associated with lower immune marker densities of common T-cell cytokines (e.g., IL17a, IL6) and NK cell ligands, consistent with an immune-colder^30^ environment relative to TAs. Here, we extend those observations by showing that TVs have significantly lower adjusted CD4⁺ and CD8⁺ mean counts compared with both TAs and SSLs. Low T-cell counts, particularly low CD4⁺ effector and CD8⁺ cytotoxic cells, are associated with reduced immune-surveillance activity, pointing to a higher likelihood for progression to CRC and consistent with the aggressive nature of the villous lesions.^13, 31, 32^ Moreover, TVs also showed reduced RRs for Th17 relative to TAs. Th17-mediated homeostatic responses play a central role in epithelial protection, barrier maintenance, and elimination of microbial products,^33, 34^ pointing to potential barrier compromise in TVs,^34^ also linked with tumor progression.^33^

The neighborhood analysis comparing TVs to TAs and SSLs highlights the emergence of an immune suppressive contexture centered around Treg and TcTreg cell types. In our prior spatial analysis^19^ comparing adenomas (TAs/TVs) to SSLs, we observed close clustering of CD4⁺ cells with Tregs, consistent with a coordinated dampening response. In the present study, TVs showed increased proximity of T-cells to Tregs and TcTregs cell types compared to SSLs and TAs. Tregs can suppress anti-tumor immunity by promoting apoptosis of cytotoxic T-cells and diminishing the effector responses.^35–37^ High abundance of Tregs are often linked with progression or poorer outcomes but in some instances may quell the proinflammatory environment, diminishing tissue damage and minimizing the chances of progression to CRC. The T-cell neighbors of Tregs seem to matter: Tregs with nearby T-cells have poorer prognosis whereas Tregs with greater proportional proximities from T-cells (like in SSLs) appear to lead to better outcomes.^15^ Recent evidence in nasopharyngeal cancer^38^ and CRC^21^ identified regions with high densities of cytotoxic T-cells clustering with Tregs as poorly prognostic. In our data, we observed that TVs had a higher proportion of reciprocal CD8^+^ to TcTreg interactions than in TAs (**Supplemental Figure 3**). TcTregs, while a rare population, are even more suppressive than Tregs.^39^ TVs also exhibited greater reciprocal Tc17-to-Treg proximity in comparison to SSLs. Tc17 cells are weakly cytolytic and, in cancer, represent a dysfunctional phenotype.^40, 41^ Together our results suggest that the greater proximities to CD8^+^, TcTregs, and Tc17 cell types in TVs may foster immune suppressive and dysfunctional neighborhoods.

TVs also demonstrated frequent proximity to “Other” (non–T-cell) cell types relative to both TAs and SSLs. T-cell cold regions are reminiscent of the immune-excluded niches of CMS3 CRCs^7, 15^ or immune deserts described for CMS2 CRCs, highlighting potential importance of underlying genetic drivers of T-cell sparseness in TVs. TVs also exhibit enhanced ECM remodeling and collagen cross-linking (e.g., elevated PLOD3 ^42^) which may physically restrict T-cell adhesion and trafficking across the TiME. TVs also show higher proliferation ^43^ (e.g., Ki-67, c-Myc, cyclin D1), suggesting high turnover and metabolic stress which may predispose T-cell exhaustion and apoptosis,^44^ in turn influencing immune neighborhoods.

### TAs as immune rich CD4 helper coordinated lesions, SSLs as low inflammation and low regulatory lesions

Contrary to expectations, TAs were CD4^+^ and CD8^+^ rich lesions overall, and more like SSLs, relative to TVs. However, neighborhood analyses showed that CD8^+^ , CD4^+^, Th17, Tc1, and Tc17 cells in TAs had closer proportional proximities to CD4^+^ cells relative to SSLs, suggesting a stronger effector response as opposed to cytotoxic. These findings are in line with prior work which consistently reports that SSLs, relative to adenomas, exhibit stronger cytotoxic signaling, whereas adenomas have higher T-helper populations.^45–47^ More recent studies further demonstrate that MSS CRCs, for which TAs and TVs are the parent lesions, have higher Th17, Treg, and Th2 cell populations than MSI-H CRCs which have higher Tc1 and Th1 responses.^14^ Moreover, spatial analysis in their work found that the interactions between Tregs, Th17, and Th2 cells in MSS CRCs clustered closer together with farther distances to Th1 and Tc1. Clustering patterns in TAs observed herein suggest that these lesions are more immunogenic than their MSS CRC counterparts. Future analysis will need to examine how TAs change in immunogenicity as they grow.

SSLs, the precursors of MSI-H CRCs, demonstrated lower Th17 cells and lower regulatory T-cell types compared to TVs. Our prior small study found that SSLs and TSAs had higher NK ligand density than conventional adenomas,^5^ and others have found that SSLs compared to conventional adenomas showed enhanced expression of cytotoxic cells (e.g., CD8⁺, NK, γδT).^45^ Intraepithelial lymphocytes and PD-1/PD-L1 densities increase with progression in the serrated pathway,^9^ consistent with epithelial dominance and a tendency toward T-cell exhaustion rather than depletion of T-cells as observed in TVs. In our previous spatial analysis study, contrasting SSLs and adenomas, we observed SSLs contained abundant CD4⁺ cells near the tumor center but Treg cells were scanty and spatially distant,^19^ consistent with present findings. A greater spatial distance between CD8^+^ T-cells and Tregs is linked to better clinical outcomes.^15^ Huang et al.^14^ reported that Th1 and Tc1 cells closely interacted within MSI-H CRCs but were spatially distant from Treg, Th2, and Th17 cells. Our NN results confirm that SSLs relative to TAs showed reduced clustering near Treg or Th17 populations. At the same time we have no evidence of strong Tc1 and Th1 spatial interactions in SSLs, as is observed in MSI-H CRCs, which could reflect the unexpectedly high immunogenicity of TAs and earlier phase of development of SSLs. It will be important to consider the spatial relationships in advanced SSLs (i.e., dysplasia) in future work.

Strengths of this study include single-cell resolution with simultaneous phenotyping and permutation-based neighborhood testing that complement bulk count summaries. Limitations include the small number of samples for TVs and SSL subtypes compared with TAs, and that we used a convenience sample from a larger cross-sectional study. In addition, the biological significance of low cell proportions in the NN analysis for TVs relative to other lesion types is difficult to interpret, given the small number of positive cells.

Future work should 1) investigate whether increasing villous component spatially correlates with reduced CD4⁺ and CD8⁺ infiltration and greater Treg clustering, as our findings suggest that spatial organization may shift with advancing villous architecture, 2) incorporate lesion size as a proxy for progression as the TiME likely changes as the lesion progresses towards malignancy, and 3) further evaluate tissue-specific (stroma, epithelium) neighborhood relationships among T-cells and “Other”. In summary, regulatory proximity, especially Treg-centered clustering, and Th17-Treg neighborhoods distinguished TVs from SSLs and TAs even when bulk T-cell counts were similar. TAs showed helper-dominant, Th17-skewed architectures, whereas SSLs appeared less effector-skewed. These spatial features refine our understanding of preinvasive colorectal immunobiology and, together with prior work, point to hypotheses that warrant testing in larger, longitudinal studies.

## Acknowledgements

This study was funded by a grant from the National Cancer Institute (R01 CA226086). Supported in part by the Biostatistics Shared Resource, Hollings Cancer Center, Medical University of South Carolina (P30 CA138313) and the Biorepository & Tissue Analysis Shared Resource, Hollings Cancer Center, Medical University of South Carolina (P30 CA138313).

