## Supplemental Tables, Figures and Methods for "Differences in T-cell counts and neighborhood patterns in human colorectal adenomas and sessile serrated lesions"

**Supplemental Material**

| Target antigen | Clone (supplier) | Dilution | Opal™ dye |
|---|---|---|---|
| **CD4** | SP35 (Roche) | predilute | 690 |
| **T-bet** | D6N8B (Cell Signaling) | 1 : 100 | 520 |
| **FoxP3** | 236A/E7 (Abcam) | 1 : 300 | 480 |
| **RORγt** | 6F3.1 (Millipore) | 1 : 100 | 620 |
| **CD8** | SP57 (Roche) | predilute | 570 |
| **Pan-keratin** | AE1/AE3/PCK26 (Roche) | predilute | 780 |

**Supplemental Table 1.** Antibody panel used for multiplex immunofluorescence (mIF) staining, including target antigen, clone (supplier), dilution, and frequencies for Opal[TM] dyes.

| Cell Type | SSL vs TA (q) | SSL vs TV (q) | TA vs TV (q) |
|---|---|---|---|
| **CD4** | 0.89 (0.43) | **1.85 (0.004)** | **2.07 (<0.001)** |
| **CD8** | 1.18 (0.39) | **1.82 (0.007)** | **1.54 (0.03)** |
| **Th1** | 0.54 (0.19) | 0.81 (0.62) | 1.49 (0.29) |
| **Th17** | 0.76 (0.39) | 1.54 (0.20) | **2.04 (0.004)** |
| **Treg** | 0.73 (0.39) | *0.41 (0.07)* | 0.56 (0.14) |
| **Tc1** | 0.77 (0.39) | 0.84 (0.62) | 1.09 (0.76) |
| **Tc17** | 0.92 (0.67) | 1.33 (0.47) | 1.44 (0.19) |
| **TcTreg** | 0.76 (0.47) | *0.36 (0.09)* | 0.47 (0.14) |

**Supplemental Table 2**. Rate ratios for T-cell type counts in SSL/TA, SSL/TV, and TA/TV comparisons adjusted for age, sex, anatomic location, size and corrected for multiple test comparisons, q < 0.05 are bolded and q < 0.10 are italicized.

| Cell Type | SSL Count (95% CI) | TA Count (95% CI) | TV Count (95% CI) |
|---|---|---|---|
| **CD4** | 21,322 (15,915-28,565) | 21,615 (18,088-25,830) | 9,460 (6,331-14,135) |
| **CD8** | 3,396 (2,491-4,629) | 2,451 (2,017-2,978) | 1,486 (952-2,320) |
| **Th1** | 211 (83-538) | 320 (126-813) | 160 (53-480) |
| **Th17** | 1,493 (1,033-2,159) | 1,757 (1,413-2,185) | 707 (442-1,131) |
| **TcTh17** | | | |
| **Treg** | 248 (150-410) | 325 (240-440) | 466 (244-889) |
| **Tc1** | 140 (70-278) | 160 (86-297) | 109 (49-240) |
| **Tc17** | 233 (154-351) | 227 (177-291) | 128 (76-215) |
| **TcTreg** | 8 (3-17) | 10 (6-15) | 15 (6-35) |

**Supplemental Table 3a**. Mean T-cell counts and 95% CIs in the stroma of SSLs, TAs, and TVs, adjusted for age, sex, anatomic location and corrected at q < 0.05.

| Cell Type | SSL Count (95% CI) | TA Count (95% CI) | TV Count (95% CI) |
| --- | --- | --- | --- |
| **CD4** | 4,837 (3,001-7,797) | 5,475 (3,502-8,561) | 2,776 (1,627-4,737) |
| **CD8** | 949 (623-1,446) | 948 (640-1,403) | 677 (413-1,110) |
| **Th1** | 34 (15-79) | 75 (34-167) | 67 (25-175) |
| **Th17** | 270 (145-501) | 370 (210-650) | 234 (117-467) |
| **TcTh17** | | | |
| **Treg** | 56 (25-127) | 64 (29-141) | 157 (60-414) |
| **Tc1** | 47 (23-93) | 62 (33-118) | 69 (30-155) |
| **Tc17** | 86 (49-150) | 82 (51-131) | 70 (37-131) |
| **TcTreg** | 3 (1-5) | 3 (2-5) | 11 (5-24) |

**Supplemental Table 3b**. Mean T-cell counts and 95% CIs in the epithelium of SSLs, TAs, and TVs, adjusted for age, sex, anatomic location and corrected at q < 0.05.

| Cell Type | SSL vs TA (q) | SSL vs TV (q) | TA vs TV (q) |
| --- | --- | --- | --- |
| **CD4** | 0.97 (0.91) | **2.45 (0.003)** | **2.54 (<0.001)** |
| **CD8** | 1.32 (0.85) | **2.47 (0.01)** | **1.88 (0.03)** |
| **Th1** | 0.66 (0.85) | 1.32 (0.54) | 2.01 (0.13) |
| **Th17** | 0.85 (0.85) | **2.11 (0.05)** | **2.49 (0.003)** |
| **Treg** | 0.76 (0.85) | 0.53 (0.23) | 0.69 (0.38) |
| **Tc1** | 0.87 (0.88) | 1.28 (0.54) | 1.47 (0.32) |
| **Tc17** | 1.02 (0.91) | 1.82 (0.19) | 1.78 (0.12) |
| **TcTreg** | 0.85 (0.88) | 0.58 (0.46) | 0.68 (0.45) |

**Supplemental Table 4a**. Rate ratios for T-cell type counts in the stroma for SSL/TA, SSL/TV, and TA/TV comparisons adjusted for age, sex, anatomic location, size and corrected for multiple test comparisons, q < 0.05 are bolded and q < 0.10 are italicized.

| Cell Type | SSL vs TA (q) | SSL vs TV (q) | TA vs TV (q) |
| --- | --- | --- | --- |
| **CD4** | 0.88 (0.72) | *1.74 (0.06)* | **1.97 (0.002)** |
| **CD8** | 1.00 (0.99) | 1.40 (0.23) | 1.39 (0.15) |
| **Th1** | *0.46 (0.06)* | 0.52 (0.22) | 1.13 (0.77) |
| **Th17** | 0.73 (0.46) | 1.15 (0.64) | 1.58 (0.13) |
| **Treg** | 0.87 (0.85) | *0.36 (0.07)* | **0.41 (0.05)** |
| **Tc1** | 0.75 (0.72) | 0.68 (0.43) | 0.90 (0.77) |
| **Tc17** | 1.05 (0.93) | 1.25 (0.58) | 1.19 (0.73) |
| **TcTreg** | 0.81 (0.85) | *0.27 (0.07)* | *0.33 (0.08)* |

**Supplemental Table 4b**. Rate ratios for T-cell type counts in the epithelium for SSL/TA, SSL/TV, and TA/TV comparisons adjusted for age, sex, anatomic location, size and corrected for multiple test comparisons, q < 0.05 are bolded and q < 0.10 are italicized.

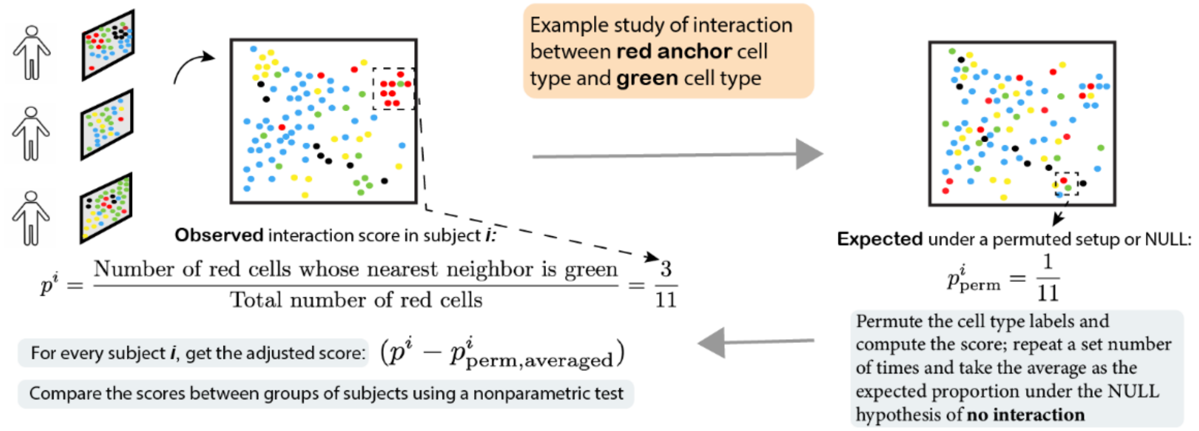


**Supplemental Figure 1:** Graphical summary of the nearest neighborhood approach.

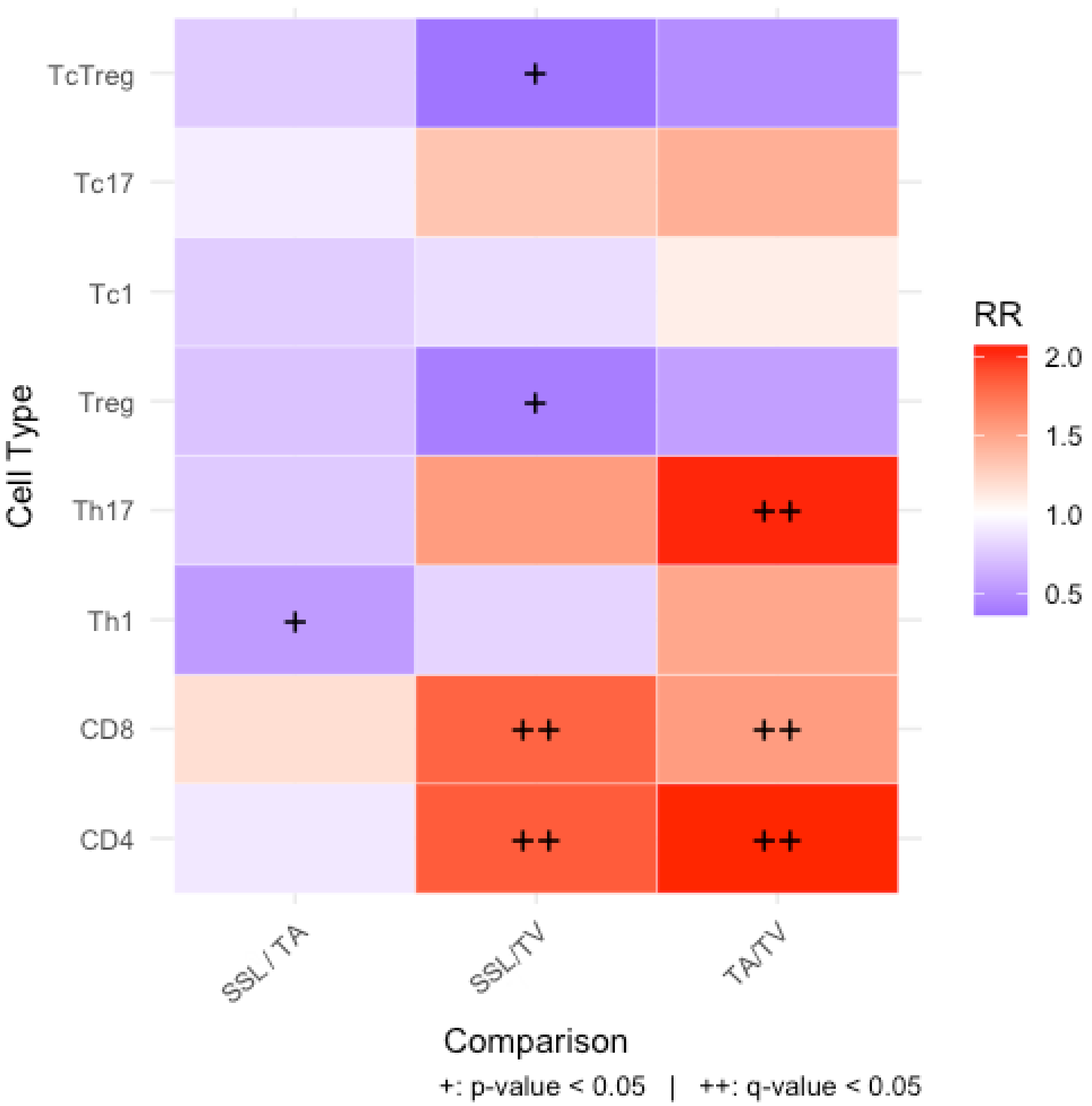


**Supplemental Figure 2**. Comparisons of T-cell rate ratios (RR) between histological types (SSL, TA, TV). Red tiles indicate greater RR for a T-cell subtype (rows) for the specified histologic type of comparison (columns). Purple tiles indicate reduced RR for a T-cell subtype for the specified histologic type comparison. Models were adjusted for age, sex, anatomic location, and lesion size.

$^{+}$p-value < 0.05

$^{++}$q-value < 0.05

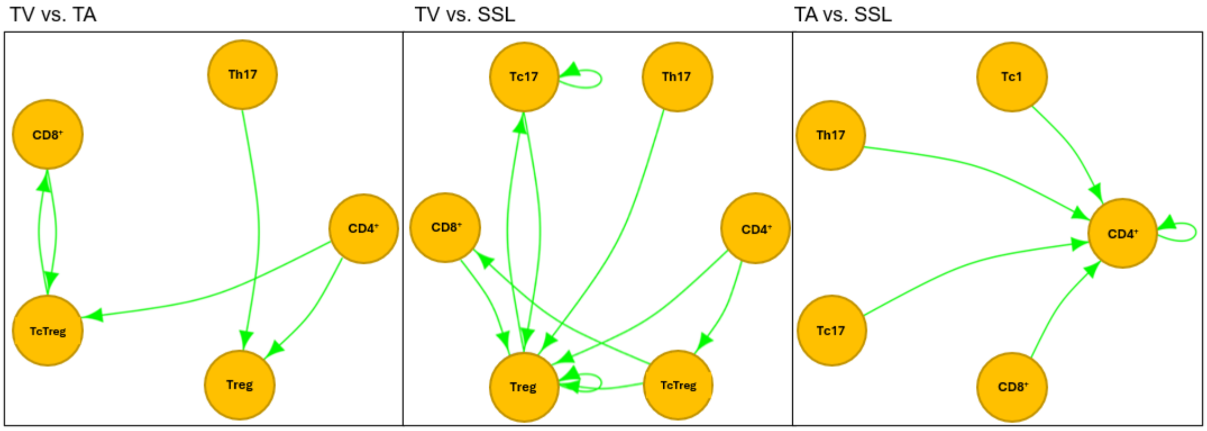


**Supplemental Figure 3**. Comparison of T-cell nearest neighbor interactions for TV vs. TA, TV vs. SSL, and TA vs. SSL. Directed arrows represent higher anchor-to-nearest neighbor T-cell type interaction in the first listed histologic type relative to the second listed histologic type (e.g., for the TV vs. TA panel (far left), the directed edges represent higher anchor-to-nearest neighbor T-cell type interaction in TVs relative to TAs.). Anchor/neighbor pairs shown are statistically significant at $q < 0.05$.

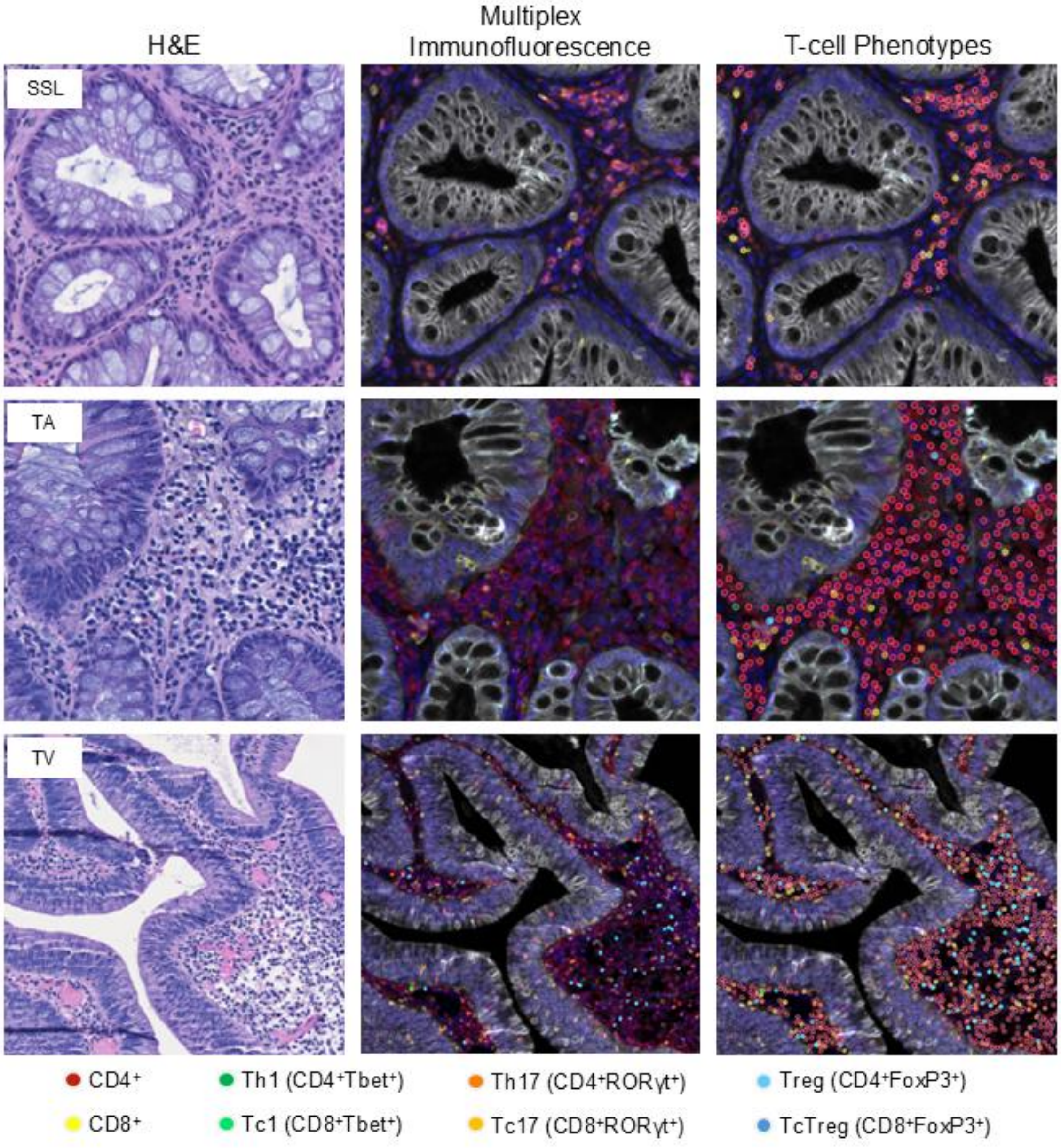


**Supplemental Figure 4**. Representative images of sessile serrated lesion (SSL, top row), tubular adenoma (TA, middle row), and tubulovillous/villous adenoma (TV, bottom row). Columns show the H&E, multiplex immunofluorescence (mIF), and T-cell phenotyping.

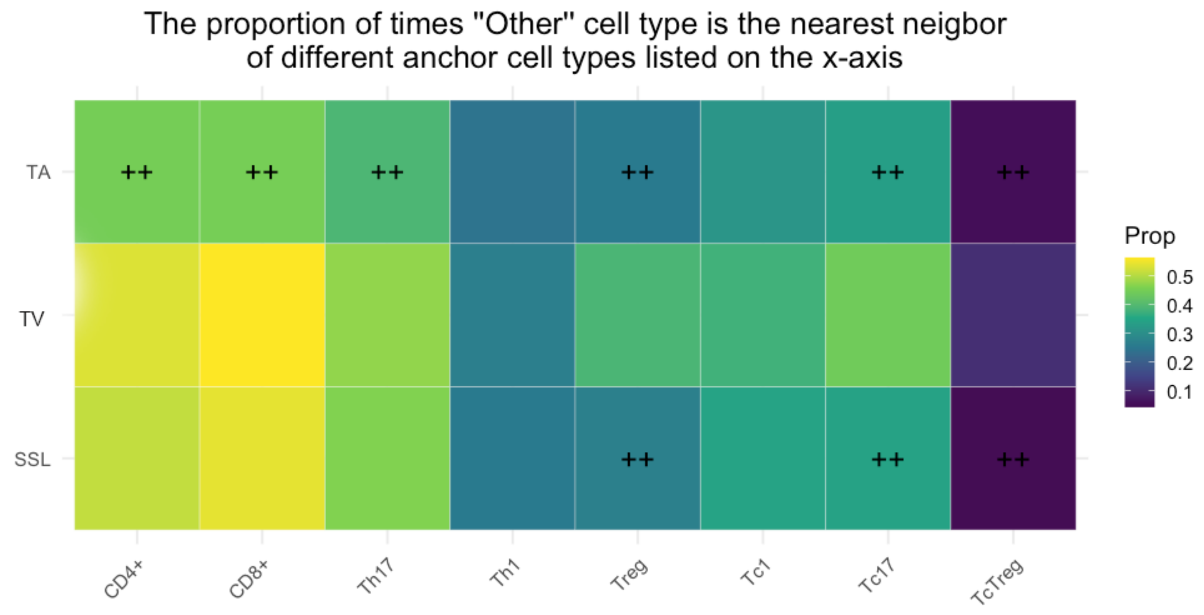


**Supplemental Figure 5**. Nearest neighbor analysis for each lesion histologic type (rows), showing the proportion of times the “Other” cell type is the nearest neighbor of each T-cell type (columns). Brighter tiles indicate a greater proportion of “Other”-T-cell neighbor occurrences.

$^{++}$(top row) significant TA/TV neighborhood comparison at $q < 0.05$

$^{++}$(bottom row) significant SSL/TV neighborhood comparison at $< 0.05$

Supplemental Methods

*Brightfield Scanning and Review*

Whole slide imaging of the H&E stained tissues was performed at 20X magnification using the Akoya Vectra® Polaris™ Automated Quantitative Pathology Imaging System (PhenoImager™ HT, Akoya Biosciences, Marlborough, MA) by the Translational Science Laboratory in the Hollings Cancer Center at the Medical University of South Carolina (Charleston, SC). Review and annotation of the whole slide scans were performed by personnel blinded to the identification of each slide using the Phenochart™ software (v[1.1.0], Akoya Biosciences, Marlborough, MA) and open-source software QuPath (v[0.2.3]).

*Multiplex Immunohistochemistry*

Polyp tissue deparaffinization and multiplex immunofluorescence (mIF) staining was performed using the Roche Ventana Discovery Ultra Automated Research Stainer (Roche Diagnostics, Indianapolis, IN). Multiplex detection was performed by pairing primary antibodies with the Akoya Opal™ multiplexing method based on Tyramide Signal Amplification for the simultaneous detection of six biomarkers (Akoya Biosciences, Marlborough, MA). The slides were stained for antibodies against: [CD4 (SP35, predilute, Roche), T-bet (D6N8B, 1:100, Cell Signaling), FoxP3 (236A/E7, 1:300, Abcam), ROR gamma T (6F3.1, 1:100, Millipore), CD8 (SP57, predilute, Roche), and Pan Keratin (AE1/AE3/PCK26, predilute, Roche)] and the fluorescence signals were generated using the following Opal™ fluorophores: [Opal 480 (FoxP3), Opal 520 (T-bet), Opal 570 (CD8), Opal 620 (ROR gamma T), Opal 690 (CD4), and Opal 780 (Pan Cytokeratin)] followed by incubation with DAPI for nuclear counterstaining. Stained slides were mounted with ProLong™ Gold Antifade Reagent (Cat. # P36934, ThermoFisher) and coverslips were applied. The mIF stained polyp tissues were imaged at 20X magnification using the Akoya Vectra® Polaris™ Automated Quantitative Pathology Imaging System (PhenoImager™ HT, Akoya Biosciences, Marlborough, MA) by the Translational Science Laboratory in the Hollings Cancer Center at the Medical University of South Carolina (Charleston, SC). Review and annotation of the whole slide scans were performed by personnel blinded to the identification of each slide using the Phenochart™ software (v[1.1.0], Akoya Biosciences, Marlborough, MA). Analysis was performed using inForm® Tissue Analysis Software (v[2.6.0], Akoya Biosciences, Marlborough, MA). Downstream analysis was performed using the PhenoptrReports Open Source R Package (https://akoyabio.github.io/phenoptrReports/index.html, Akoya Biosciences, Marlborough, MA).

Image analysis of mIF-stained tissues was performed using inForm® Tissue Analysis Software (v[2.6.0], Akoya Biosciences, Marlborough, MA). The workflow included Tissue Segmentation, Cell Segmentation, and Phenotyping. For training across these workflow features, at least 10 regions of interest were selected among the lesions for analysis. Lesions for training varied in histologic type to best capture the biological and morphological heterogeneity across lesion types. The regions of interest were used in the following steps:

Tissue Segmentation: Tissue regions (epithelium, stroma) were defined using the *Trainable Tissue Segmentation* feature. For tissue segmentation training, specifications were required for Tissue Categories, Components for Training, Pattern Scale, and Segmentation Options. Three Tissue Categories were defined: Epithelium, Stroma, and Other (denoting non-tissue regions). DAPI (nuclear marker) and pan-keratin (epithelial marker) were selected as Components for Training, that is, only DAPI and pan-keratin and no other biomarkers were considered for defining tissue regions. A Large pattern scale was selected, as suggested by the inForm User Guide v.3.3.0. for tissue sections and larger scale structures. For smoothing edges along tissue regions, a Medium segmentation resolution was selected and Trim Edges was set to 5 pixels in Stroma regions within the Segmentation Options.Multiple iterations of training for tissue segmentation were implemented to ensure adequate performance of the segmentation algorithm across morphologically heterogeneous samples.

Cell Segmentation: The *Segment Cells* feature was used for cell segmentation, including segmentation of the nuclei, cytoplasm, and cell membrane. Additional specifications were required for Nuclear Component Splitting, Assisted Component Splitting, Other Settings, and Membrane and Cytoplasm Settings. Under Components, DAPI was selected for nuclear detection and pan-keratin was selected to aid in membrane detection and nuclear splitting, given this marker is primarily expressed along the cell membrane. Specifications in Nuclear Component Splitting included a mixture of quality for Nuclear Staining, a Splitting Sensitivity of 0.33, and a Minimum Nuclear Size of 40 pixels. For Assisted Component Splitting, the Assisting Staining was "a mixture of quality," a Splitting Sensitivity of 0.40, and a Minimum Nuclear Size of 35 pixels. Other Settings indicated to fill nuclear holes smaller than 2 pixels and to refine cells after segmentation. Lastly, the following specifications were selected for Membrane and Cytoplasm Settings: a Cytoplasm Thickness of 3 pixels, a Membrane Search Distance of 10 pixels, a mixture of quality for Membrane Staining, and a Membrane Signal Threshold of 0.50.

Cell Phenotyping: *Phenotyping* was performed for the six markers of interest: CD4, CD8, pan-keratin, T-bet, RORγt, and FoxP3. Together, these would indicate the following T-cell phenotypes: CD4 ($CD4^+$), Th1 ($CD4^+$T-$bet^+$), Th17 ($CD4^+$$ROR\gamma t^+$), Treg ($CD4^+$$FoxP3^+$), CD8 ($CD8^+$), Tc1 ($CD8^+$$Tbet^+$), Tc17 ($CD8^+$$ROR\gamma t^+$), and TcTreg ($CD8^+$$FoxP3^+$). Additional dual T-cell subtypes were defined using individual cell biomarker positivities [e.g., Th17-Treg ($CD4^+$$ROR\gamma t^+$$FoxP3^+$)], as indicated in the main text. In the Phenotyping Settings, a Phenotyping Schema was created for each biomarker and were set to a Standard Associated View. For each of the six biomarkers, cells were indicated as positive for the biomarker (e.g., FoxP3+ for cells positive for FoxP3) or "Other" for cells negative for the specified biomarker. To train this phenotyping classifier, at least 5 but no more than 40 cells were manually indicated as either positive or negative for each biomarker.
Downstream analysis was performed using the PhenoptrReports Open Source R Package (https://akoyabio.github.io/phenoptrReports/index.html, Akoya Biosciences, Marlborough, MA).
